# Innate fear responses are reflected in the blood epigenome of rhesus macaques

**DOI:** 10.1101/2020.11.05.369538

**Authors:** Hector Bravo-Rivera, Roy Lardenoije, Dimaris Merced, Adaris Mas-Rivera, James E. Ayala, Joana P. Gonçalves, Antonia V. Seligowski, Tanja Jovanovic, Kerry J. Ressler, Torsten Klengel, Gregory J. Quirk

## Abstract

Fear and anxiety are complex physiological states aimed at promoting adaptive behaviors. They are also core symptoms of many neuropsychiatric disorders; yet, our knowledge of the underlying biological correlates remains fragmented. Non-human primate models are critical for our understanding of mechanisms associated with complex higher-order behavioral phenotypes. Here we investigated individual variations in innate fear responses to a snake stimulus in free-ranging rhesus macaques and discovered an unusual bimodal distribution of fearful and fearless behavior, likely as a result of an environmental insult by a hurricane. In a translational approach, we discovered a DNA methylation profile associated with fear behavior in these monkeys. We also found evidence that this epigenetic signature is associated with innate fear responses in humans in the form of acoustic startle. Our data highlight the importance and translational utility of non-human primate models for neuropsychiatric research and provide a potential epigenetic signature of innate fear.

## 1. Introduction

Defensive fear responses to predators are fundamental for the survival of a species. In the wild, fear and avoidance responses often conflict with foraging for food, which is equally essential. It is well established that monkeys reared either in the wild or in captivity avoid snakes or snake-like objects^1–4^. However, studies have shown that some monkeys can overcome this fear behavior when the presence of snakes coincides with the presence of food^5^. This suggests that there may be considerable variability in monkeys’ fear responses to snakes, especially when fear behavior conflicts with foraging. Prior data indicate that fear responses across species share hard-wired circuits for phylogenetically relevant stimuli^6^. However, little is known about the biological factors associated with the behavioral variability observed in non-human primates, the influence of the environment, and how these are related to fear responses in humans. Epigenetic modifications such as DNA methylation have been associated with fear- and stress-related phenotypes in pre-clinical and clinical studies^7–13^ and can provide a mechanism for the interaction of genetic and environmental factors to determine behavioral outcomes^14,15^.

To identify biological and non-biological factors that may be associated with fearful reactions toward snakes, we conducted experiments on free-ranging rhesus macaques (*Macaca mulatta*) at Cayo Santiago, part of the Caribbean Primate Research Center (CPRC), a 38-acre island off the coast of Humacao, Puerto Rico, which is the birthplace of ~9 generations of rhesus macaques. Free-ranging monkeys offer the advantage of studying innate fear responses without the influence of captivity-induced stressors^16^. Previous studies at Cayo Santiago have identified biomarkers associated with behavioral traits such as social status^17–19^ and reproductive behavior^20–22^. The center’s database provides access to birth/death dates and pedigree data, making the Cayo Santiago an ideal environment to evaluate fear behavior in a natural setting.

To assess innate fear responses in these monkeys, we developed a snake-induced startle task in which a monkey must open a food bin with a hinged lid and reach past an unexpected rubber snake to retrieve desirable fruit (grape). Our goal was to identify environmental, and epigenetic factors that might correlate with variability in the monkeys’ fear responses. Our results provide evidence for a bimodal distribution of the fear response in older monkeys, which is related to the occurrence of Hurricane Georges in 1998. In addition, we detected an epigenetic DNA methylation profile that was associated with fear response. Further, we showed an overlap of epigenetic profiles with innate fear responses in humans, highlighting the importance of translational non-human primate research for understanding complex neuropsychiatric behaviors.

## 2. Results

### 2.1. Variability in fear responses to the snake stimulus

In the presence of a randomly selected monkey, a grape was placed in an otherwise empty bin of the experimental apparatus and the monkey was allowed to retrieve the grape (**Figure 1A**). The apparatus was then rotated, and a grape was placed in a bin containing a concealed snake attached to the lid. Upon opening the bin, most monkeys dropped the lid quickly when the snake appeared and refused to retrieve the grape. However, we observed a range of responses with some monkeys holding onto the lid and retrieving the grape (**Video 1**). We therefore calculated the ratio of lid holding time (with snake/without snake) as a measure of the monkey’s fearlessness (**Figure 1B**). In most monkeys, fearlessness ratios were < 1 (n = 139, 81%), indicating a fearful response to the snake stimulus. However, 32 monkeys (19%) showed fearlessness ratios ≥ 1, indicating low fear of the snake stimulus. We found that this distribution was bimodal (*b* = 0.80) and therefore classified monkeys with a fearlessness ratio < 1 as fearful and those ≥ 1 as fearless.

**Figure 1.**
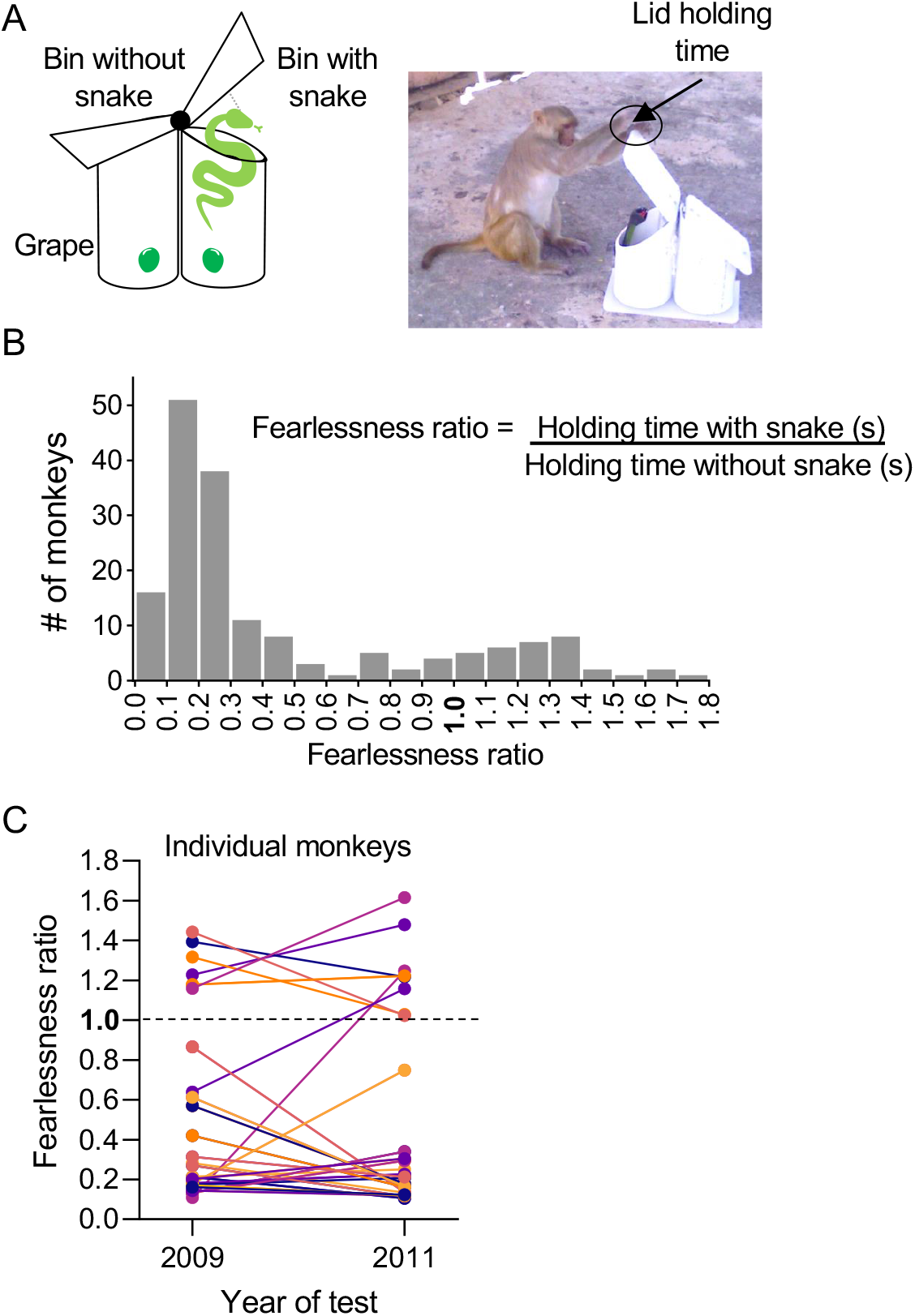
Variable fear responses to the snake stimulus. **(A)** Experimental design: We placed a seedless grape in a bin with a hinged lid allowing the monkey to lift the lid and retrieve the grape. A rubber snake was hanging from the lid so that the monkey would be exposed to the snake upon lifting the lid. **(B)** We calculated the ratio of lid-holding times (with snake/without snake) to measure the monkey’s fear response. For most monkeys, holding ratios were < 1 (n = 139), reflecting fear of the snake. However, 32 monkeys showed holding ratios ≥ 1, indicating the absence of fear to the snake. (**C**) We retested a subset of monkeys with the snake two years after the initial test with the snake and compared fear responses. We found that 6 out of 6 fearless monkeys remained fearless at retest, and 32 out of 34 fearful monkeys remained fearful.

### 2.2. Fear reactions were specific to the snake stimulus

The snake stimulus likely triggered fear responses because it resembled a live snake. Alternatively, the novelty and unexpectedness of a hanging object could trigger fear responses^23^. To address these questions, we substituted the snake with a novel object (Nerf ball) in a subset of monkeys who had never been exposed to the snake (**Supplementary Figure 1**). These monkeys showed significantly less fear to the ball than to the snake (mean fearlessness ratio for ball = 1.23; mean fearlessness ratio for snake = 0.40; t_(180)_ = 3.65; p < 0.001), corroborating that fear reactions to the snake were not due to novelty or unexpectedness.

### 2.3. Snake reactions were consistent across time

To test if fearlessness ratios were consistent within monkeys, we retested a subset of monkeys two years after the initial test (**Figure 1C**). We found that 32 out of 34 monkeys initially classified as fearful remained fearful at retest, whereas 6 out of 6 monkeys initially classified as fearless remained fearless. Furthermore, there was no significant difference in the average fearlessness ratios at the two time points (2009 = 0.52; 2011 = 0.40; t_(39)_ = 0.23, p = 0.82). This suggests that fear reactions to the snake stimulus are persistent and resemble a trait.

### 2.4. Fearlessness is associated with age, coinciding with hurricane exposure

We observed a positive correlation between fearlessness and age (r_Pearson_ = 0.428, p < 0.001) (**Figure 2A**) and fearless monkeys were significantly older than fearful monkeys (fearful = 10.3 years, fearless = 16.23 years, t_(170)_ = 7.35; p < 0.01). A notable increase in the incidence of fearlessness is apparent in the age range of 10-13 years. This suggests that age may interact with biological and/or environmental factors to modulate responses to the snake. Both male monkeys (n = 111) and female monkeys (n = 60) showed the same relationship between age and fearlessness ratio (**Supplementary Figure 2A and 2B**), and there was no significant differences between sexes (male = 0.45; female = 0.47; t_(170)_ = 0.36; p = 0.72).

**Figure 2.**
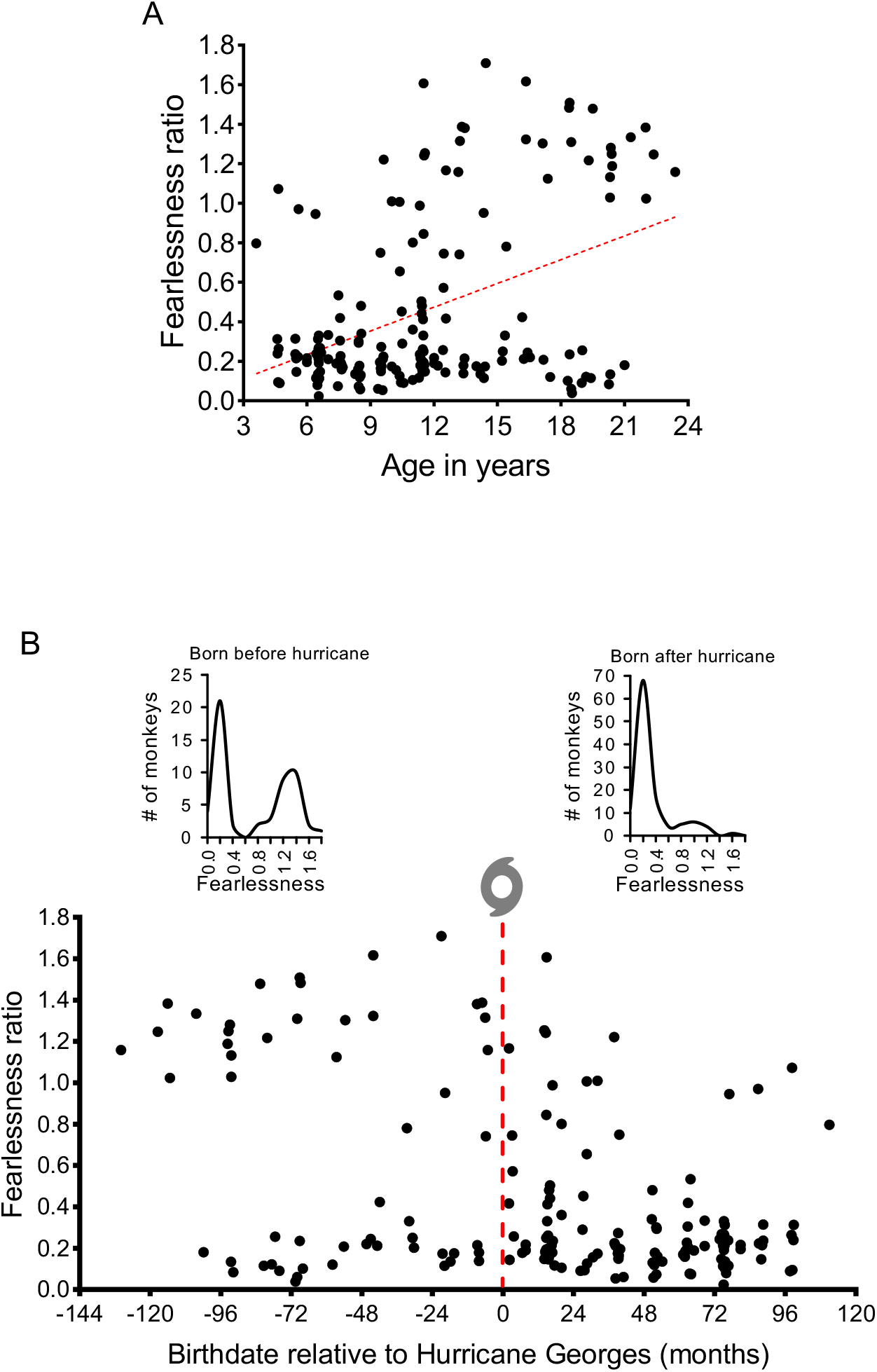
Fear responses to the snake correlate with age and prior exposure to a hurricane. **(A)** Fearlessness was significantly correlated with age (r_Pearson_ = 0.42; p < 0.001), and fearless monkeys were significantly older than fearful monkeys. **(B)** Monkeys born before Hurricane Georges divided into distinct fearless and fearful subgroups. In contrast, monkeys born after were mostly fearful of the snake.

To better understand the adaptive (or maladaptive) nature of our fearlessness measure, we used reproductive fitness to assess evolutionary success, and correlated fearlessness ratios with the number of male monkeys’ offspring. (**Supplementary Figure 2C**). For each monkey, the number of offspring was divided by the monkey’s reproductive age (actual age – 4 years, the start of reproductive behavior)^24^. Fearlessness was positively correlated with reproductive fitness (r_Pearson_ = 0.32; p < 0.001), suggesting that fearless monkeys might have an evolutionary advantage in mating. This measure of reproductive fitness does not apply to female monkeys because they are generally limited to a single offspring per mating season.

To gain insight into the potential heritability of fearlessness, we asked whether genetically similar monkeys showed similar fear responses. For this, we used the kinship coefficient (KC), a measure of the expected proportion of genetic similarities between two individuals. KC data were acquired from the CPRC database and were calculated using several variables (pedigree, sex, birth date)^20^. Comparison of each pairs’ fearlessness ratio and their KC did not yield a significant correlation (r_Pearson_ = 0.006; p = 0.45), suggesting that variance in this fear response is not aggregated in families.

The age range at which fearlessness increased (10-13 years) does not correspond with any known biological change in adult monkeys (e.g. puberty), and suggests an effect of experience. Interestingly, 10-13 years prior to each monkey’s test corresponds to the occurrence of a major Category 4 hurricane that made landfall on Cayo Santiago in 1998: Hurricane Georges^25^. Re-plotting the monkeys’ birthdates relative to the occurrence of Hurricane Georges (**Figure 2B**) shows that monkeys born prior to the hurricane were divided into distinct fearless and fearful subgroups. In contrast, monkeys born after the hurricane were mostly fearful. While correlational, this suggests that a history of hurricane exposure may have reduced fear responses to the snake in a subset of the older group (24/54) of monkeys.

### 2.6. Hurricane-exposed monkeys showed distinct DNA methylation patterns associated with fearlessness

To investigate the association between the fear response and DNA methylation, we adapted the Illumina Infinium MethylationEPIC array platform, designed to assess methylation states across the human genome^26^, for use with rhesus macaque DNA samples (**Figure 3A**; see Methods section for details). Briefly, the probe sequences were mapped to the rhesus macaque genome, and 343,441 high-confidence human-macaque probes remained after exclusion of probes with a mismatch in the 5 positions closest to the target site (see Methods section). Epigenome-wide association studies (EWAS) were performed on a total of 88 monkeys (27 hurricane-exposed and 61 non-exposed). Models were fitted with the fearlessness ratio as the main predictor, and age, sex, as well as 1 surrogate variable (SV, to control for unobserved confounders) as covariates for the exposed and non-exposed animals separately (**Supplementary Figures 4A, B**).

**Figure 3.**
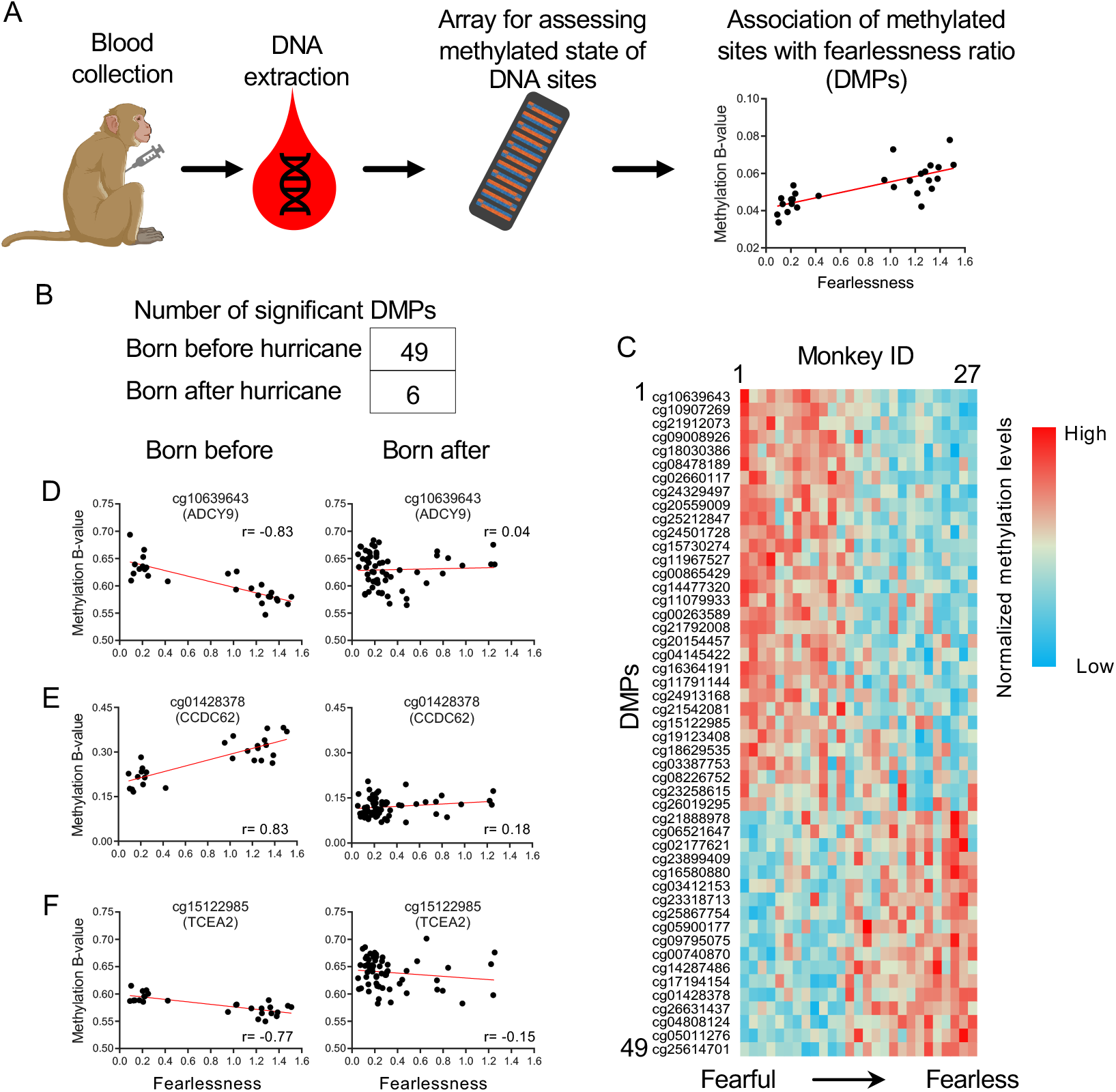
Association of fearlessness ratio with the methylation state of specific DNA sites. **(A)** Blood samples were collected from monkeys, and DNA was extracted, bisulfite converted, and processed on Infinium MethylationEPIC arrays to investigate genome-wide DNA methylation levels. **(B)** We observed 49 differentially methylated positions (DMPs) associated with fearlessness in monkeys born before the hurricane (n = 27), compared to 6 such DMPs in monkeys born after the hurricane (n = 61). **(C)** Heatmap of normalized methylation values for the monkeys born before the hurricane. Columns represent individual monkeys sorted from left to right in ascending order of fearlessness ratio (fearful to fearless). Rows represent each of the 49 DMPs correlated with fearlessness, showing their normalized methylation value adjusted for covariates (individual beta value/average of beta values for that DMP). **(D-F)** Methylation patterns for 3 representative DMPs (see Discussion) from monkeys exposed (left) or non-exposed (right) to Hurricane Georges. Each plot shows the beta values of that site (with 0 indicating no methylation and 1 full methylation) vs. the fearlessness ratio, Pearson r- and (FDR-adjusted) p-values result of linear regression analysis between fearlessness and β-values.

The EWAS identified 49 differentially methylated positions (DMPs, individual methylation sites) associated with fearlessness in the hurricane-exposed animals (**Figure 3**; **Table 1**; **Supplementary Table 1**); 18 (37%) showed a positive association of methylation with fearlessness, whereas 31 (63%) showed a negative association with fearlessness (**Figure 3C**). In the non-exposed animals, 6 DMPs were identified, of which 3 (50%) showed a positive association with fearlessness and 3 (50%) showed a negative association (**Supplementary Figure 3**; **Table 2**). There was no overlap in DMPs between the exposed and non-exposed animals. This was followed by a regional analysis, which identified 15 differentially methylated regions (DMRs, a group of methylation sites; 8 hypermethylated, 7 hypomethylated) associated with fearlessness in the monkeys that experienced the hurricane (**Table 3**), and (3 hypermethylated) in those that did not (**Table 4**). Between the two groups there was no overlap in genes with a DMR. Thus, we detected fearlessness-associated differential DNA methylation mostly in hurricane-exposed monkeys.

**Table 1.** Top 10 differentially methylated positions (DMPs) associated with fearlessness in monkeys exposed to Hurricane Georges. Shown for each DMP is the associated site ID, associated genomic position (Mmul_10), relative location within a CpG island (CGI), closest gene in the rhesus macaque genome (ncbiRefSeq), gene feature (CDS, coding sequence), log2 fold change (FC), mean M value, degrees of freedom (DF), t-value, p-value, and false discovery rate-adjusted p-value (q-value).

| Site | Position | CGI feature | Closest gene | Gene feature | log2FC | Mean | DF | t | p | q |
| --- | --- | --- | --- | --- | --- | --- | --- | --- | --- | --- |
| cg01428378 | chr11:122634961 |  | CCDC62 | Intron | 0.74 | -2.76 | 22 | 6.351 | 2.14E-10 | 7.34E-05 |
| cg26019295 | chr11:53617209 | shore | HOXC6 | Intergenic | -0.26 | -3.61 | 22 | -6.116 | 9.59E-10 | 1.65E-04 |
| cg25614701 | chr16:51464308 |  | RARA | Intergenic | 0.23 | -6.06 | 22 | 5.856 | 4.74E-09 | 5.42E-04 |
| cg10639643 | chr20:4059314 |  | ADCY9 | Intron | -0.31 | 0.67 | 22 | -5.774 | 7.73E-09 | 6.13E-04 |
| cg18030386 | chr19:4876366 | shore | UHRF1 | Intron | -0.54 | 3.72 | 22 | -5.75 | 8.92E-09 | 6.13E-04 |
| cg05011276 | chr20:23462911 | shore | LCMT1 | Intergenic | 0.45 | -4.21 | 22 | 5.711 | 1.12E-08 | 6.43E-04 |
| cg16364191 | chr4:42048385 | shore | NHSL1 | CDS | -0.53 | 1.98 | 22 | -5.632 | 1.78E-08 | 7.89E-04 |
| cg10907269 | chr7:164226179 |  | DYNC1H1 | Intron | -0.4 | 4.45 | 22 | -5.627 | 1.84E-08 | 7.89E-04 |
| cg11079933 | chr9:104427391 |  | LOC114670143 | Intergenic | -0.41 | 1.37 | 22 | -5.549 | 2.88E-08 | 1.10E-03 |
| cg04808124 | chr4:51812443 |  | SHPRH | Intergenic | 0.25 | -5.06 | 22 | 5.456 | 4.87E-08 | 1.67E-03 |

**Table 2.**
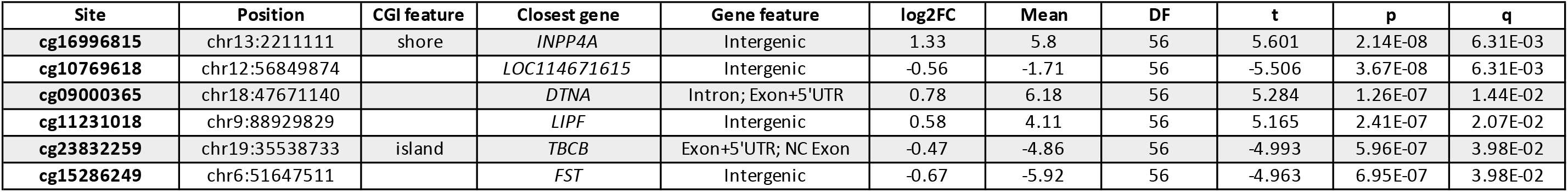
Differentially methylated positions (DMPs) associated with fearlessness in monkeys not exposed to Hurricane Georges. Shown for each DMP is the associated site ID, associated genomic position (Mmul_10), relative location within a CpG island (CGI), the closest gene in the rhesus macaque genome (ncbiRefSeq), gene feature (NC, non-coding; UTR, untranslated region), log2 fold change (FC), mean M value, degrees of freedom (DF), t-value, p-value, and false discovery rate-adjusted p-value (q-value).

**Table 3.**
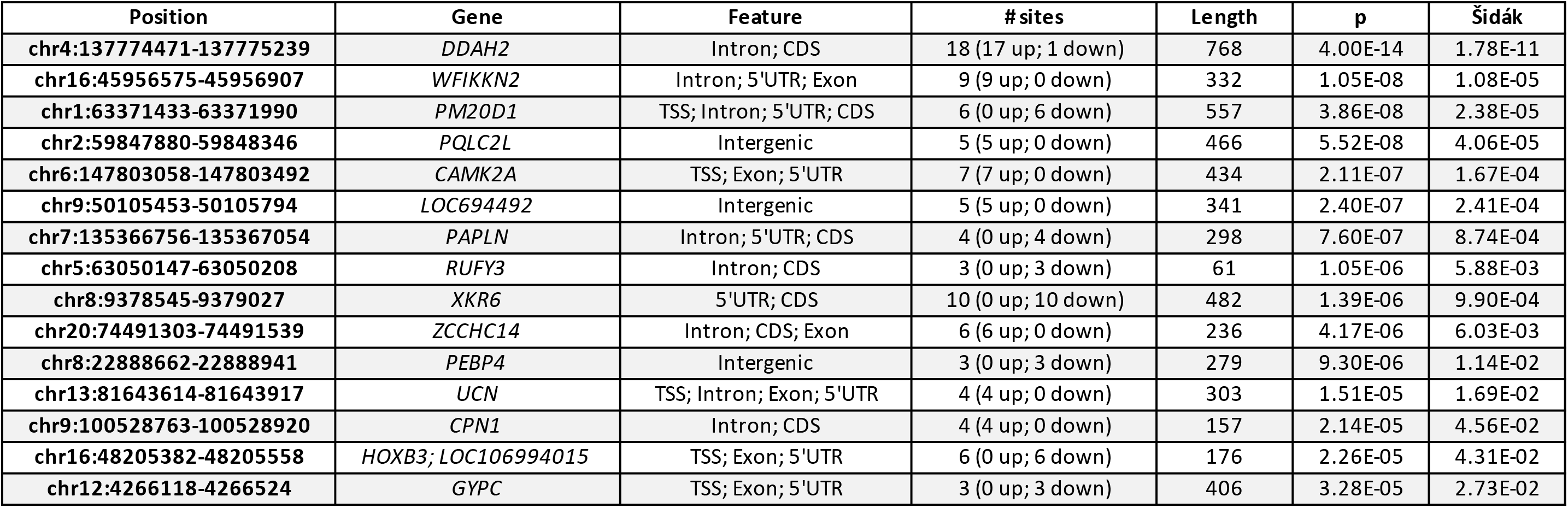
Differentially methylated regions (DMRs) associated with fearlessness in monkeys exposed to Hurricane Georges. Shown for each DMR is the genomic position (Mmul_10), closest gene (ncbiRefSeq), gene feature (CDS, coding sequence; TSS, transcription start site; UTR, untranslated region), number of sites constituting the region (and number of up/down regulated sites), length of the region (in bp), p-value, and Šidák-adjusted p-value.

**Table 4.** Differentially methylated regions (DMRs) associated with fearlessness in monkeys not exposed to Hurricane Georges. Shown for each DMR is the genomic position (Mmul_10), closest gene (ncbiRefSeq), gene feature (CDS, coding sequence), number of sites constituting the region (and number of up/down regulated sites), length of the region (in bp), p-value, and Šidák-adjusted p-value.

| Position | Gene | Feature | #sites | Length | p | Šidák |
| --- | --- | --- | --- | --- | --- | --- |
| chr4:137936929-137936979 | *TNF* | Intron; CDS | 4 (4 up; 0 down) | 50 | 3.32E-08 | 2.28E-04 |
| chr7:92960291-92960422 | *COCH* | Intergenic | 3 (2 up; 1 down) | 131 | 9.42E-07 | 2.46E-03 |
| chr14:108750236-108750476 | *LOC114672549* | Intergenic | 4 (4 up; 0 down) | 240 | 1.15E-06 | 1.64E-03 |

### 2.7. Fearlessness-associated sites in monkeys rank higher in association with startle response in humans

In order to investigate a potential link between the DNA methylation signature of fearlessness in rhesus macaques and fear-related phenotypes in humans, we leveraged a separate collection of MethylationEPIC array and startle psychophysiology data from the Grady Trauma Project, in which acoustic startle responses were associated with DNA methylation levels (see Methods section for details). After similar pre-processing as the monkey data, 780,559 probes remained from 134 trauma-exposed subjects with acoustic startle response data. A total of 46 of the 49 fearlessness-associated sites found in the hurricane-exposed monkeys were present in the processed human data. EWAS of the human data with the startle response (electromyogram amplitude - EMG) as a predictor did not yield genome-wide significant results, which was expected given the overall sample size. Of note, effect sizes for the monkey EWAS (log2 fold change range −0.79 to 0.74) were much larger than those in the human analysis (log2 fold change range −0.0013 to 0.0012). Nevertheless, the 46 sites associated with fearlessness in the monkeys had significantly lower p-values compared to all other sites in the human EWAS (Wilcoxon rank-sum test p-value = 3.03E-04; permutation p-value = 3.00E-04; **Figure 4**), suggesting that the DNA methylation signature of fearlessness in rhesus macaques may be relevant for the startle response in humans.

**Figure 4.**
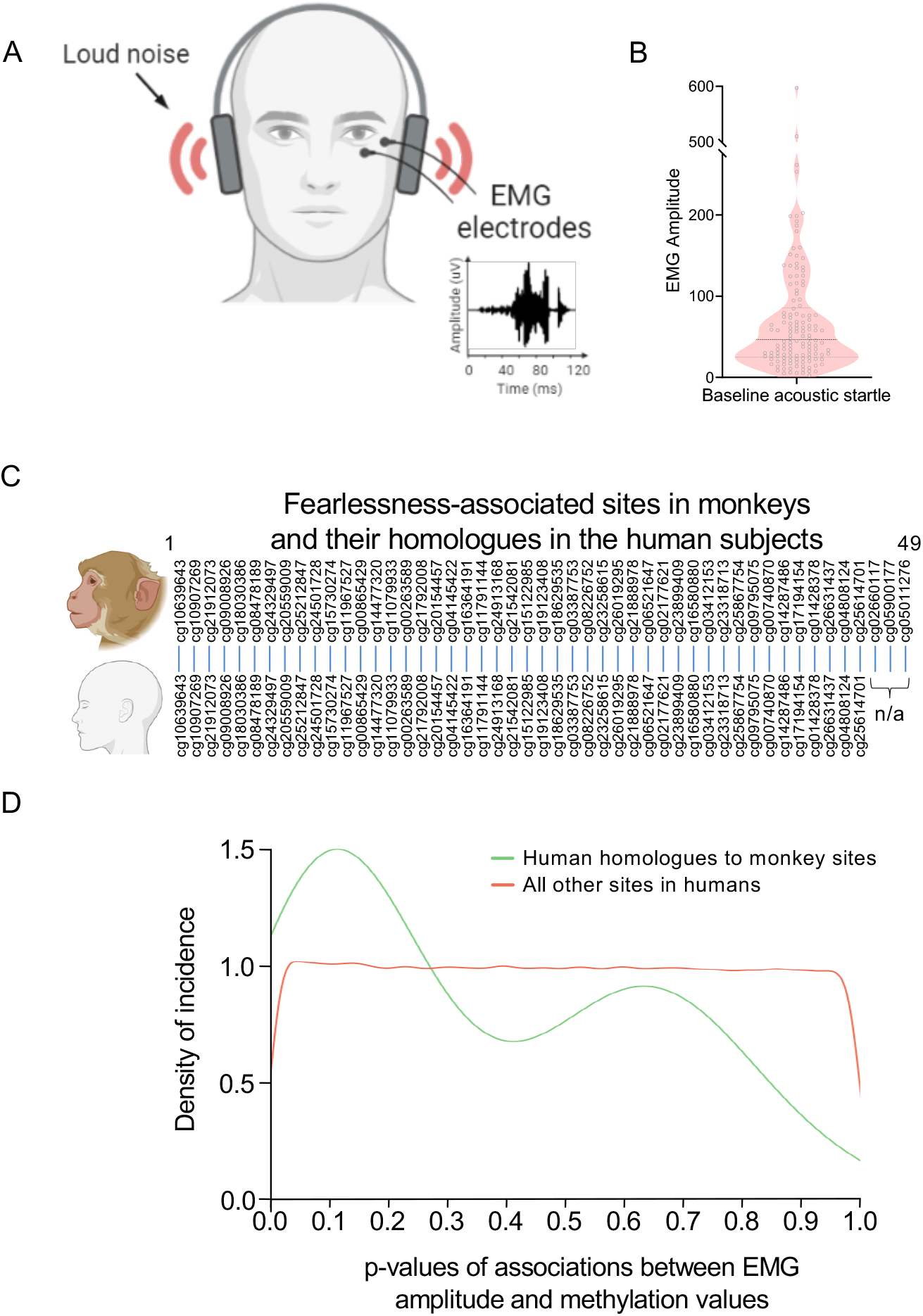
In humans, homologues of monkey fearlessness sites show increased association with innate startle responses. **(A)** Human subjects were presented with an unexpected loud noise (108db). EMG recordings from the orbicularis oculi record startle responses. **(B)** Distribution of baseline acoustic startle responses across 104 subjects (defined as maximum amplitude of the orbicularis oculi muscle contraction 20 to 200ms following the startle probe) **(C)** Out of 49 fearlessness associated sites in monkeys, we identified 46 homologues in humans. **(D)** Densities of the p-values from human startle and epigenome association, for the sites shown in C (n=46) compared with all other sites (n = 780,513). A Wilcoxon rank sum test comparing these two sets of p-values indicates that the fearlessness sites have significantly lower p-values (p = 3.03E-04), as is reflected in the plot by the density peak at the lower p-values for the fearlessness sites.

## 3. Discussion

We developed a simple behavioral task in free-ranging rhesus macaques to assess fear responses to a snake stimulus in a natural environment. Prior studies demonstrated that wild-reared or laboratory raised rhesus macaques show strong fear to a snake stimulus, albeit in the latter, this response seems less pronounced^16,27–29^. In most studies, the latency to reach over a rubber snake to acquire food is the central phenotype^28,30–32^. This approach allows monkeys to observe the snake stimulus for a period of time and evaluate the risks involved in obtaining the food reward. The latency to reach for the reward may therefore be determined by the monkeys’ assessment of the risk, rather than by their initial fear reaction. In our task, the exposure to the snake is unexpected, as revealed by a short lid holding time (i.e. dropping the lid of the box when suddenly exposed to the snake), similar to the startle reflex to an unexpected stimulus.

In the 171 animals we examined, older monkeys were more likely to be fearless of the snake than younger individuals. Increased reproductive success in fearless animals suggests that this trait correlates with other adaptive behaviors. Prior studies showed that a minority of wild reared monkeys were not afraid of snakes^2,16,27^. We corroborate these findings and provide additional evidence suggesting that rhesus macaques show considerable differences in the immediate response to the snake stimulus. In fact, our data revealed an age-dependent bimodal distribution of fear reactions, which is in contrast to observations that most behavioral phenotypes follow a single Gaussian distribution^33^. We concluded that a bimodal distribution with no overlap indicates the presence of one or more factors dividing the sample into distinct subgroups. Although rare, previous studies have observed bimodal distributions of conditioned fear responses without overlap in rats and monkeys^16,34^.

We are not aware of significant developmental changes at the age range of 10-13 that would justify such a salient behavioral change^35^. It is conceivable that monkeys’ prior experience with snakes on the island could have affected their fear responses in our task. However, unlike predatory snakes usually found in natural habitats of rhesus macaque in central and southeast Asia^36,37^, *Alsophis portoricens,* the species of snake found on Cayo Santiago typically feeds on lizards and is not known to attack monkeys. Furthermore, it has been suggested that possible observation of other monkeys’ testing by a conspecific may result in observational conditioning^2^. While we cannot rule out possible effects of either prior snake encounters or observational conditioning, we think it is unlikely that such encounters would result in a non-overlapping bimodal distribution of fearlessness. Also, retest stability across a 2-year time frame indicates that potential confounders or stressors coinciding with the time of testing, such as interactions with a more dominant monkey, are an unlikely driver of this phenotype.

Interestingly, the emergence of this distinct phenotype in older individuals coincided with the devastating effects of a hurricane in the region. Evidence from human clinical studies and animal models including non-human primates, suggests that traumatic experiences can lead to compensatory adaptations in behavior^38–41^. While it seems an appealing hypothesis that Hurricane Georges may be causal to the increase in fearlessness we observed, our opportunistic study cannot prove causality and disentangle the effects of age and hurricane exposure. However, irrespective of the true underlying cause, we hypothesized that the divergent fear responses may be represented in biological correlates detectable in these animals.

Epigenetic modifications, broadly defined, regulate gene expression without a change in the DNA sequence. A broad body of evidence suggests that environmental and experiential factors can trigger cascades of epigenetic modification that in turn mediate downstream molecular and behavioral phenotypes. In this study, we focused on DNA methylation in rhesus macaque and human blood samples using the flexibility of Illumina DNA methylation arrays as a translational platform to investigate DNA methylation in both human and non-human primate species^42–45^. In fact, we were able to identify 49 DMPs and 15 DMRs in the hurricane-exposed animals after correction for multiple testing. Although the functional implications of the identified DMPs/DMRs remain unknown, some of the associated genes have been previously implicated in fear-related phenotypes or have potential functional implications not yet explored. For example, *ADCY9* encodes the ubiquitously expressed adenylyl cyclase-9 implying a role in beta-adrenergic modulation of sympathetic responses^46^. *TCEA2* has recently been associated with trait anxiety in the largest GWAS of anxiety to date from the Million Veterans Program^47^. Moreover, a DMR in *PM20D1* similar to our results was recently identified in human blood samples in a longitudinal study of post-traumatic stress disorder^48^, and *CAMK2A* is one of the most well-understood genes mediating synaptic plasticity underlying fear learning in a variety of paradigms^49^.

Undoubtedly, the significance of peripheral epigenetic signals for central nervous system functions remains controversial and requires further work^50,51^. Since non-human primates and humans share neuronal circuits and mechanisms that drive innate behaviors such as startle responses, we sought to corroborate our results across species. We found that DMPs associated with the monkeys’ fear responses show a stronger association than other sites with startle responses in humans^52^. Our study provides evidence that the detection of DNA methylation profiles in non-human primates living in semi-controlled environments may accelerate epigenetic approaches in human studies confounded by additional environmental factors. At the same time, our study exemplifies the potential of environmental factors such as natural disasters to shift behavioral and epigenetic traits, thereby improving our understanding of humans exposed to stressful or traumatic environments.

## 4. Methods

### 4.1. Study site and subjects

The study was conducted out on Cayo Santiago, a 0.14 km^2^ island located 1 km off the east coast of Puerto Rico, which is inhabited by ~1000 free-ranging rhesus macaques (*Macaca mulatta*), whose ancestors were brought from India in the 1930s ^53^. At Cayo Santiago, monkeys are provided with food and water, and yearly blood draws are performed as part of the veterinary care. Our research was approved by the Institutional Animal Care and Use Committee of the University of Puerto Rico (protocol #A3340109). Subjects consisted of 171 adult monkeys, 4 - 25 years of age, with 111 males and 60 females. Monkeys were obtained from seven social groups located in different parts of the island: group F (n = 57), group HH (n = 27), group S (n = 9), group KK (n = 16), group R (n = 37), group V (n = 19), and group X (n = 6). Monkeys on Cayo Santiago are provisioned daily with commercial monkey chow and water and can browse natural vegetation. Data were collected over a two-year period (2009 - 2011), May through September, during the hours of 7:00 AM - 12:00 PM.

### 4.2. Behavioral Data

We constructed a plastic testing apparatus consisting of two identical PVC bins with hinged lids, which could be opened to retrieve a grape. A 35 cm long rubber snake was hung from the underside of one of the lids with a nylon fishing line (**Figure 1A**, **Video 1**). Subjects were encountered randomly across the island. Monkeys had been previously identified via a tattoo on the chest and inner left thigh. We enticed subjects with grapes to encourage participation. After a monkey showed interest, we placed a grape in the safe bin (no snake) in the monkeys’ presence and allowed the monkey to lift the lid and retrieve the grape. This process occurred twice. The third grape was placed in the bin that contained the rubber snake attached to the lid. Monkeys opened the lid and responded to the snake when it appeared. All grape retrievals were videoed from a distance of approximately 4 m. To quantify fear responses, we compared the amount of time the subjects held the lid in trials with and without the snake: 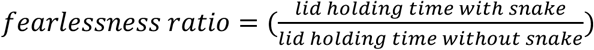. Only data from the first trial (for both no-snake and snake) were used. Because monkeys tended to drop the lid faster on snake trials, fear responses to the snake yielded a holding ratio < 1, while fearless responses yielded a ratio ≥ 1. The bimodality coefficient was given by the formula 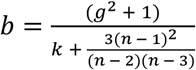, where *n* is the number of monkeys, *g* is the sample skewness, and *k* is the sample excess kurtosis. Values greater than 0.55 indicate a bimodal distribution.

### 4.3. Video analysis

To assess the amount of time spent holding the lid, we identified the video frame in which the monkey’s hand contacted the lid and the video frame where the hand terminated contact with the lid (video rate was 30 frames/second). Subtracting these frame counts and dividing by 30 yielded the holding time in seconds. The holding time was calculated for all monkeys (n = 171) for trials with and without the snake. In a subset of monkeys, we assessed the height of the lid opening (n = 49) and the orientation of the snake (facing toward or away from the monkey, n = 17).

### 4.4. Blood collection

A subset of the monkeys in our study was captured for blood draws (n = 147). Capture dates ranged from January to March in both 2010 and 2012. Monkeys were lured with food into a feeding corral approximately 100 m^2^, which was provisioned daily with commercial monkey chow. Once inside the corral, monkeys were captured by net, then transferred into a cage and transported to a sampling station located on the island. Trapping occurred between 8:30 AM - 12:00 PM. All subjects were inspected by a veterinarian and found to be in generally good health. Subjects were anesthetized with ketamine (10 mg/kg, i.m.), and blood samples were collected between 7:30 AM - 10:30 AM. Subjects captured after 10:30 AM remained in the cage until the next day. Whole blood was collected in 5 ml EDTA tubes and placed into a cooler containing ice packs. Upon returning to the mainland (at 12:00 PM), tubes were frozen at −80°C until extraction at AKESOgen Inc (Peachtree Corners, GA) using the Gentra Puregene Blood Kit (Qiagen). DNA concentration was determined using pico green. A total of 500 ng DNA was then bisulfite converted using the EZ-96 DNA Methylation Kit (Zymo) according to the instructions of the manufacturer, hybridized to Infinium MethylationEPIC microarrays (Illumina), and scanned using an iScan (Illumina), according to the instructions of the manufacturer, at AKESogen Inc. Raw data in the form of IDAT files from a total of n = 90 animals with overlapping blood and phenotype information was exported and subsequently processed in R as described below.

### 4.5. Pedigree data

We obtained pedigree data from the CPRC long-term database. This database contains maternal assignments based on census information such as the behaviors of the putative mothers (e.g., lactation) for all individuals from the founding population onwards and maternity and paternity microsatellite markers for most monkeys born since 1990 (approximately 2900 monkeys). From our sample, maternity was based on 48 subjects, and paternity was based on 87 subjects. There was a 97.4% agreement between the census data and genetic data in the population as a whole. Missing paternity links can result in underestimation of additive genetic variance; however, paternity error rates < 20% introduce few biases in genetic analyses^54^. There is no evidence of high rates of inbreeding on Cayo Santiago^55^.

### 4.6. Probe mapping and annotation

To determine which MethylationEPIC probes could be used, the procedure outlined by Needhamsen and colleagues^56^ for mice was adapted for rhesus macaques. In short, the MethylationEPIC probe source sequences were extracted from the MethylationEPIC manifest file provided by Illumina (v1.0 B4; https://support.illumina.com/array/array_kits/infinium-methylationepic-beadchip-kit/downloads.html) and converted to the FASTA format using the *write.fasta* function of the seqinr R package (version 3.4-5)^57^, and the rhesus macaque genome (Mmul_10) was downloaded from Ensembl (release 98)^58^. Bismark (version 0.17.0) was then used for *in silico* bisulfite conversion of the macaque genome and mapping the MethylationEPIC probe source sequences to the converted genome with a seed sequence of 28 nucleotides and allowing only unique hits with up to 2 mismatches (-l 28 -n 2)^59^. Using the SAMtools (version 1.4.1)^60^ calmd (-e) function, true mismatches were detected as inconsistencies between C > T conversions in MZ tags and mismatches in MD tags of Bismark-generated BAM files. Only probes with no mismatches within 5 bp of the target site were selected, and probes mapping to non-standard chromosomes were dropped. The remaining sites were annotated with the Mmul_10 “refGene” and “cpgIslandExt” University of California, Santa Cruz (UCSC) genome browser tables using CruzDB (version 0.5.6)^61^.

### 4.7. DNA methylation data analysis

DNA methylation data analysis was performed with R (version 3.6.2) and RStudio Pro (version 1.2.5033)^62^ unless otherwise specified. Data files are accessible through GEO GSE159027 (https://www.ncbi.nlm.nih.gov/geo/query/acc.cgi?acc=GSE159027, a reviewer password can be obtained via the editor). Raw IDAT scanner files were imported into R using the *readEPIC* function of the wateRmelon package (version 1.30.0)^63^. Only probes that were successfully mapped to the macaque genome were included. Predictions based on X chromosome methylation were performed with the *predictSex* function of the wateRmelon package and indicated no sex mismatches, after which the 10,544 probes targeting the X and Y chromosomes were removed from further analysis. Bisulfite conversion rates were computed with the *bscon* wateRmelon function, and one sample with a bisulfite conversion rate < 90% was excluded. Samples and probes were further filtered by detection p-value and beadcount using the *pfilter* function from wateRmelon, excluding samples with > 5% of sites with a detection p-value > 0.05 and sites with beadcount < 3 in > 5% of samples or a detection p-value > 0.05 in > 5% of samples. Finally, outlying samples based on a principal component analysis were detected using the *outlyx* function of wateRmelon and removed. These steps resulted in the removal of 22,878 probes. The data were then normalized using quantile normalization plus beta-mixture quantile normalization, as recommended by Wang et al^64^. Normalization violence was inspected using the *qual* function of the wateRmelon package.

The presence of genetic variants can interfere with probe binding and thereby cause extreme values. However, reliable databases for genetic variants in rhesus macaques are not available. To counter this, probe-wise outlier detection and removal was performed with the *pwod* wateRmelon function, using a threshold of 4 interquartile ranges. To limit the number of missing values introduced by this procedure, they were imputed using the k-nearest neighbors algorithm (with k = 100). Samples with > 10% and probes with > 20% missing values were excluded, resulting in the exclusion of 347 additional probes. Batch effects (array barcode, position on array, and transfer plate) were removed using ComBat^65^ without explicitly conserving the effects of interest to prevent the potential downstream inflation of test statistics^66^. After processing, the data was visually inspected for abnormalities, and one additional sample with an abnormal density plot was excluded. Since M values have better statistical properties than beta values when fitting linear models^67^, any 0 or 1 beta values were replaced with the lowest > 0 or highest < 1 beta value, respectively, after which the beta values were converted to M values using the *beta2m* function of the lumi package (version 2.38.0)^68^.

Samples from hurricane-exposed and non-exposed animals were then split for further analysis. To investigate the impact of fearlessness on DNA methylation, a model with M values as the outcome, fearlessness ratio as a predictor, and age and sex as covariates was then used for a surrogate variable analysis (SVA) with the sva package (version 3.34.0)^69^. This analysis was done to detect and capture unobserved sources of variation, such as potential differences in cell-type composition, into surrogate variables^65,70^. The number of SVs to be included in the model was determined through a proportion of variance explained plot. The model was then fit for each probe using limma (version 3.42.2)^71^, followed by an adjustment of the test statistics for bias and inflation with the bacon package (version 1.14.0)^72^. P-values were corrected for multiple testing using the false discovery rate (FDR) procedure (q-value), and DMPs were defined as sites with q < 0.05.

Following the site-wise analysis, spatial correlations between p-values were investigated with comb-p (version 0.50.1)^73^ to detect DMRs. Comb-p, a Python-based command-line tool, was run with a seeding p-value of 0.05, a window size of 1000 bp, and using Mmul_10 for annotating the regions. DMRs were defined as regions with at least 3 sites, and a Šidák corrected p-value < 0.05.

### 4.8. Human DNA methylation and startle response comparison

Although fearlessness, as assessed for this study, has not been investigated in humans, the reaction to the snake in the experiment is akin to a startle response because it is innate and reflexive, and startle responses to different stimuli have been investigated in humans. To get a first impression of how a fearlessness DNA methylation profile in monkeys might be related to similar behavior in humans, we investigated this profile in relation to the acoustic startle response, using human blood MethylationEPIC data (n = 795) from the GTP (http://gradytraumaproject.com/). This data has been described previously^74^ (GEO accession number GSE132203). The human data were analyzed similarly to the monkey data, with some minor adjustments. Due to the availability of human SNP reference data, the probe-wise outlier detection step was skipped, and potentially cross-hybridizing probes and probes close to SNPs with a major allele frequency > 0.05 in any population were directly removed^75^. The MethylationEPIC arrays for male and female GTP samples were processed in separate batches. Therefore, in addition to array barcode and position on array, the sex batch effect was also removed with ComBat. As with the monkey data, after processing, only trauma-exposed samples were selected. An individual was defined as being exposed to childhood or adult trauma based on a combination of published cut-off values for the Childhood Trauma Questionnaire (CTQ)^76^ and the presence of one or more adult trauma types as assessed by the Traumatic Events Inventory (TEI)^77^: CTQ sexual abuse subscale > 8, CTQ physical abuse subscale > 8, CTQ physical neglect > 8, CTQ emotional abuse > 10, CTQ emotional neglect > 15, and/or TEI total types experienced > 0.

Eyeblink startle data within the GTP cohort were derived from the habituation phase of a fear-potentiated startle (FPS) paradigm^78–81^. During habituation, 4 trial of 108 dB, 40 ms duration startle probes were delivered through headphones to assess baseline startle. Startle data were collected with Biopac MP150 for Windows (Biopac Systems, Inc., Aero Camino, CA). The eyeblink startle response was measured using electromyography (EMG) of the right orbicularis oculi muscle. All impedances were less than 6 kΩ. EMG activity was acquired at a sampling rate of 1 kHz, amplified, and digitized using the EMG module of the Biopac system. Startle response variables were derived from raw Biopac EMG recordings using MindWare (MindWare Technologies, Inc). Startle response was identified as the maximum amplitude of the eyeblink muscle contraction 20 to 200 ms following the startle probe.

The association between the startle response and DNA methylation was then analyzed for each site, as was done for the monkeys. Apart from the main startle predictor, age was included as a covariate, together with estimated blood cell type proportions of CD8T cells, CD4 cells, natural killer (NK) cells, B cells, and monocytes, as well as the first genotype principal component. Blood cell type proportion estimation and genotyping for the GTP cohort have been previously reported in ^74^ and ^82^, respectively. Since these covariables could be included in the model directly for the human data, SVA was not used to estimate them.

To investigate the general association of the fearlessness-associated sites with the startle response in humans, a one-sided Wilcoxon rank-sum test was used, using the p-values from the human startle EWAS to rank the sites and the monkey fearlessness DMPs to split them (i.e., it was tested if the monkey fearlessness DMPs generally had lower p-values than the other sites in the human EWAS analysis). To confirm the significance of the rank-sum test p-value, the test was repeated 100,000 times with a number of randomly sampled sites, equal to the number of fearlessness DMPs. The proportion of p-values lower than the original Wilcoxon test was determined.

## Acknowledgment statement

The authors would like to thank Aranza Torrado-Tapias, Enmanuelle Pardilla, Adrian Rivera, Oscar Ortiz, and Amie Madrigal for assistance with the behavioral experiments on Cayo Santiago. We thank Angelina Ruiz-Lambidez, Julio Resto, Giselle Caraballo-Cruz, Nahiri Rivera, and the staff members of Cayo Santiago and the Caribbean Primate Research Center who have contributed to census data collection and provided technical/administrative support. Prof. Jeffrey Rogers, Baylor College of Medicine, consulted on the kinship analysis. JPG received funding from National Institutes of Health U54EY032442, Convergence for Health and Technology 50.05.19, and HollandPTC-Varian. AVS received funding from the American Heart Association #20CDA35310031. TJ received funding from NIMH R21MH098212 and NIMH R01MH110364. KJR was supported by NIMH P50MH115874, NIMH R01MH108665 and by the Connor Group Kids and Community Partners fund. GJQ received support from NIMH R37MH058883 and NIMH P50MH106435. TK received support from ERA-NET Neuron 01EW2003, The Brain and Behavior Foundation (NARSAD) YI 20895, NICHD R21HD088931, NICHD R21HD097524, and NIMH R21MH117609. Cayo Santiago is supported by the Office of Research Infrastructure Programs (ORIP) of the National Institute of Health, grant numbers 8P40OD012217 and 3P40OD012217-27S1 awarded to the University of Puerto Rico Medical Sciences Campus, PI: Melween I. Martínez. The publication’s content is the authors’ sole responsibility and does not necessarily represent the official views of National Center for Research Resources (NCRR), Office of Research Infrastructure Programs (ORIP), or the University of Puerto Rico.

## 5. Figure and Table Legends

### Supplemental Figures

**Supplemental Figure 1.**
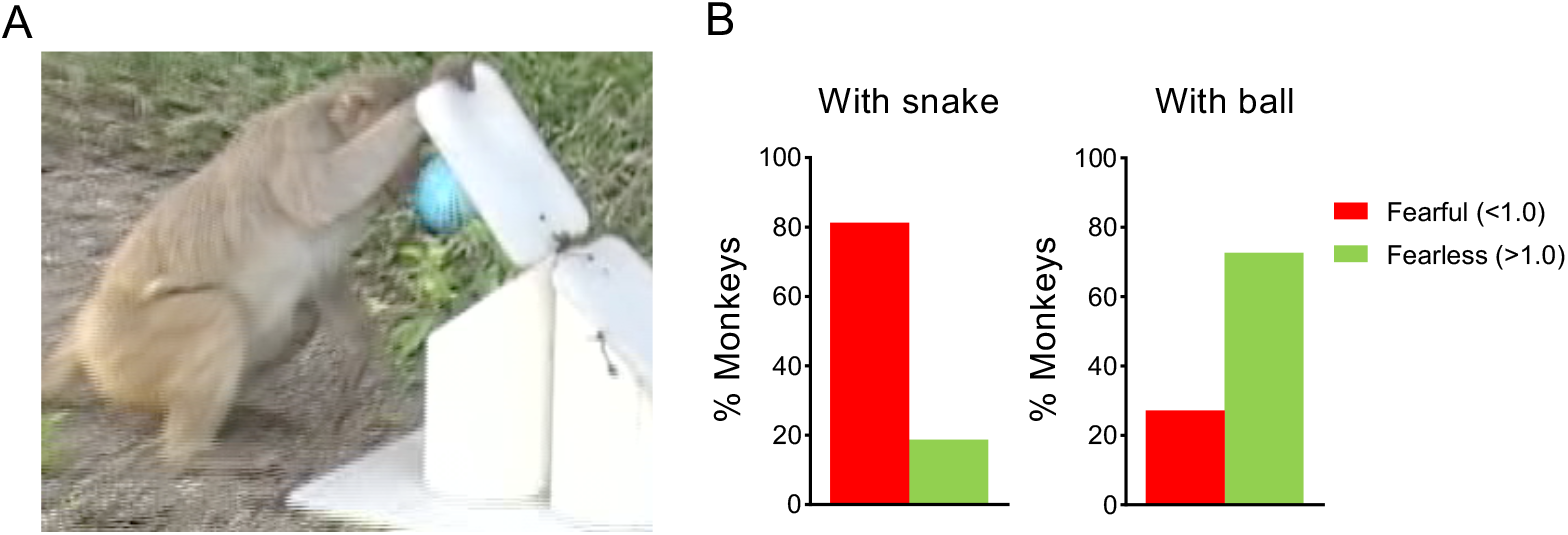
(For the main Figure 1) Fear reactions were specific to the snake stimulus. **(A)** We substituted the rubber snake with a different object (Nerf ball) in a subset of monkeys who had never been exposed to the snake. **(B)** Monkeys showed significantly less fear to the ball than to the snake (mean holding ratio for the ball: 1.23; mean holding ratio for snake: 0.40; t_(180)_ = 3.65; p < 0.001), corroborating that fear reactions to the snake were not due to novelty.

**Supplemental Figure 2.**
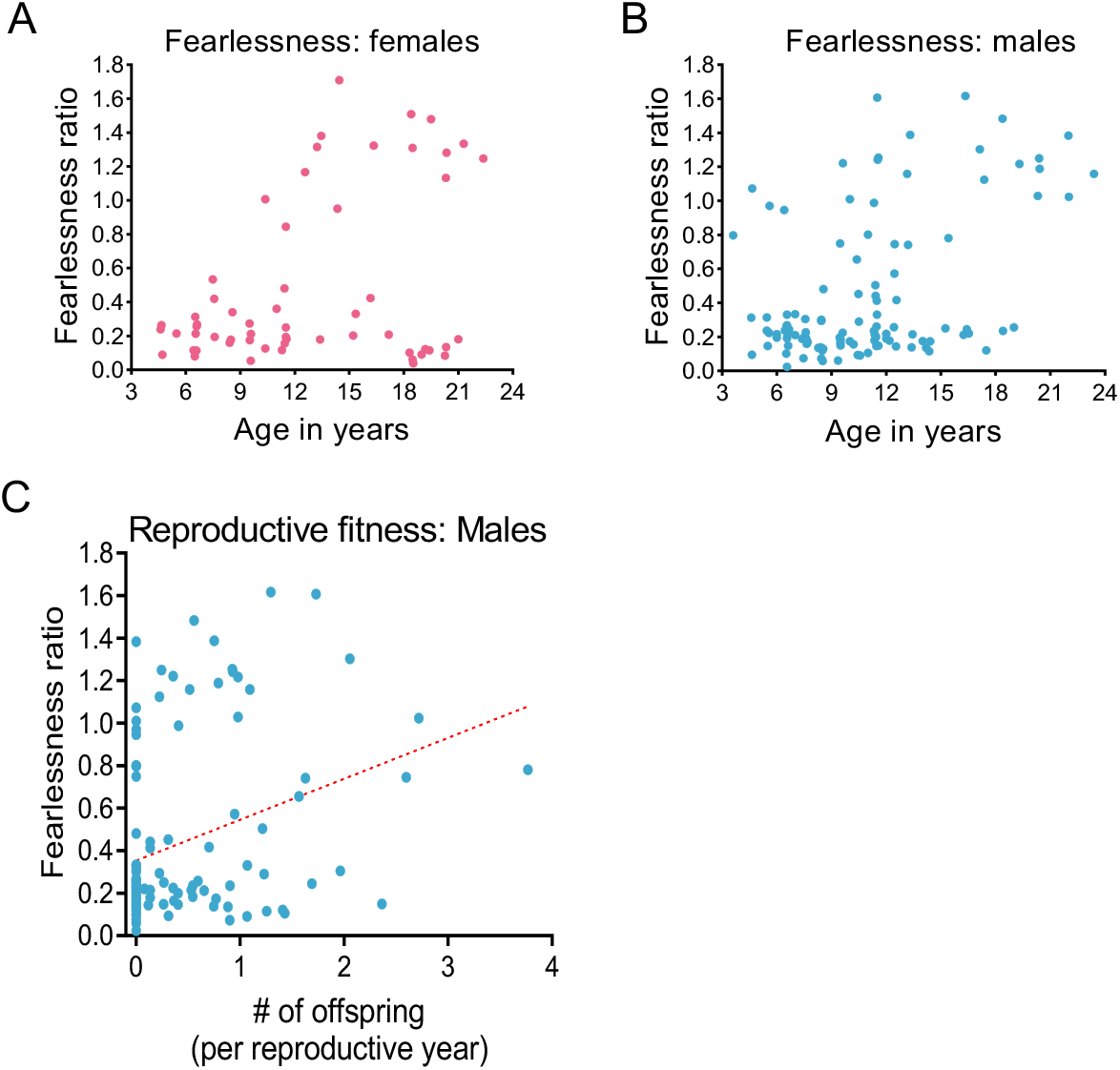
(For main Figure 2) Fearlessness is not associated with sex, but correlates with reproductive fitness. **(A-B)** Both sexes showed the same relationship between age and fearlessness ratio. There was no significant sex difference in fearlessness ratios across all ages (males mean: 0.45; females mean: 0.47; t_(170)_ = 0.36; p = 0.72), < 12 years of age (p = 0.19), or ≥ 12 years of age (p = 0.57). **(C)** Fearlessness ratio was positively correlated (r_Pearson_ = 0.32; p < 0.005) with the number of offspring in male monkeys, suggesting that fearless monkeys might have an evolutionary advantage.

**Supplementary Figure 3.**
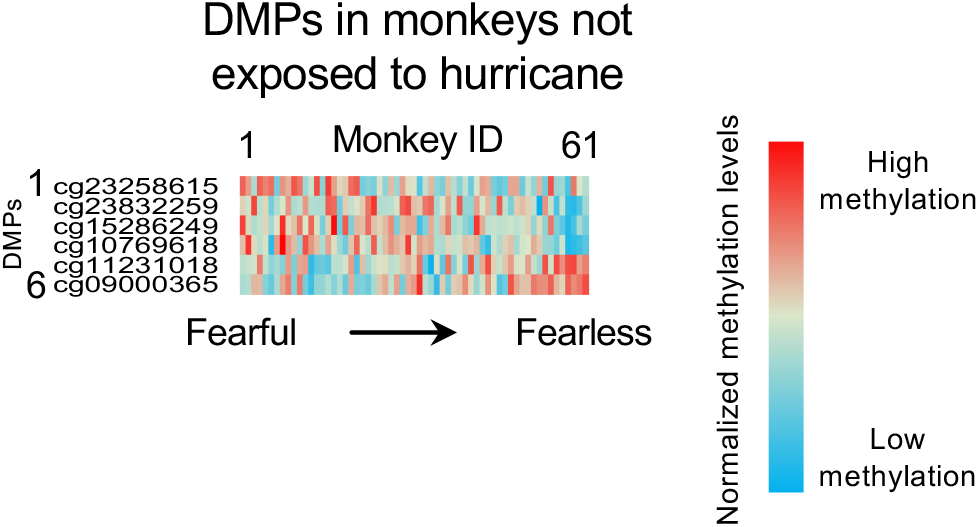
(For main Figure 4) Association of fearlessness ratio with the methylated state of specific DNA sites in monkeys not exposed to the hurricane. Heatmap of normalized methylation values for the monkeys born after the hurricane. Columns represent individual monkeys sorted from left to right in ascending order of fearlessness ratio (fearful to fearless). Rows represent each of the 6 DMPs correlated with fearlessness, showing their normalized methylation value (individual beta value/average of beta values for that DMP).

**Supplemental Figure 4.**
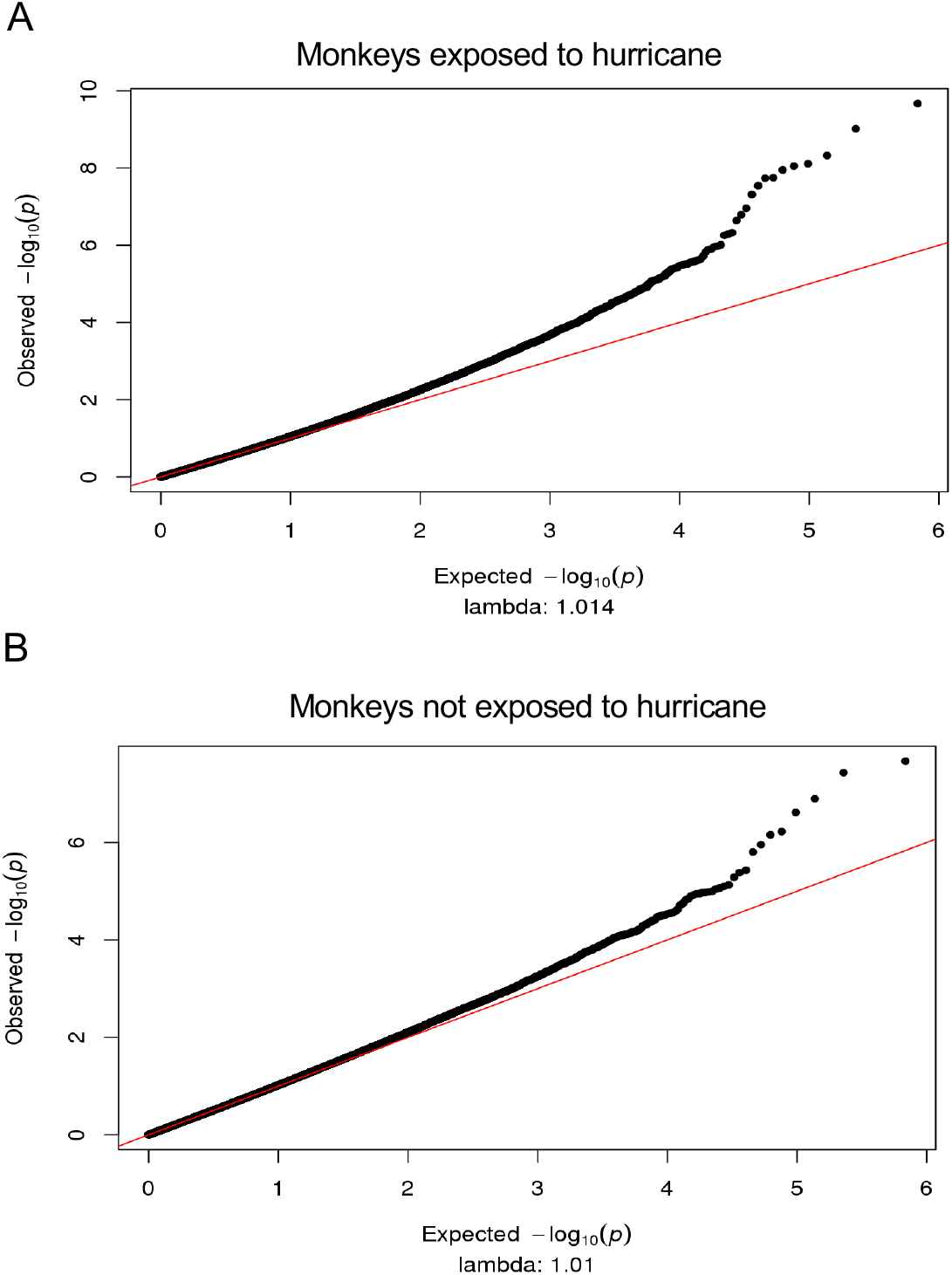
Quantile-quantile plots of the p-values from the fearlessness ratio epigenome-wide association studies in the hurricane-exposed (**A**) and non-exposed (**B**) monkeys.

### Supplemental Tables

**Supplemental Table 1.** Full list of methylated positions (DMPs) associated with fearlessness in monkeys exposed to Hurricane Georges (n = 49). Shown for each DMP is the associated site ID, associated genomic position (Mmul_10), relative location within a CpG island (CGI), closest gene in the rhesus macaque genome (ncbiRefSeq), gene feature (CDS, coding sequence), log2 fold change (FC), mean M value, degrees of freedom (DF), t-value, p-value, and false discovery rate-adjusted p-value (q-value).

| Site | Position | CGI feature | Closest gene | Gene feature | log2FC | Mean | DF | t | p | q |
| --- | --- | --- | --- | --- | --- | --- | --- | --- | --- | --- |
| cg01428378 | chr11:122634961 |  | CCDC62 | intron | 0.74 | -2.76 | 22 | 6.351 | 2.14E-10 | 7.34E-05 |
| cg26019295 | chr11:53617209 | shore | HOXC6 | intergenic | -0.26 | -3.61 | 22 | -6.116 | 9.59E-10 | 1.65E-04 |
| cg25614701 | chr16:51464308 |  | RARA | intergenic | 0.23 | -6.06 | 22 | 5.856 | 4.74E-09 | 5.42E-04 |
| cg10639643 | chr20:4059314 |  | ADCY9 | intron | -0.31 | 0.67 | 22 | -5.774 | 7.73E-09 | 6.13E-04 |
| cg18030386 | chr19:4876366 | shore | UHRF1 | intron | -0.54 | 3.72 | 22 | -5.75 | 8.92E-09 | 6.13E-04 |
| cg05011276 | chr20:23462911 | shore | LCMT1 | intergenic | 0.45 | -4.21 | 22 | 5.711 | 1.12E-08 | 6.43E-04 |
| cg16364191 | chr4:42048385 | shore | NHSL1 | cds | -0.53 | 1.98 | 22 | -5.632 | 1.78E-08 | 7.89E-04 |
| cg10907269 | chr7:164226179 |  | DYNC1H1 | intron | -0.4 | 4.45 | 22 | -5.627 | 1.84E-08 | 7.89E-04 |
| cg11079933 | chr9:104427391 |  | LOC114670143 | intergenic | -0.41 | 1.37 | 22 | -5.549 | 2.88E-08 | 1.10E-03 |
| cg04808124 | chr4:51812443 |  | SHPRH | intergenic | 0.25 | -5.06 | 22 | 5.456 | 4.87E-08 | 1.67E-03 |
| cg21912073 | chr15:67716949 |  | CCDC171 | intron | -0.35 | 1.32 | 22 | -5.308 | 1.11E-07 | 3.45E-03 |
| cg23318713 | chr9:63891429 | island | PLAU | intergenic | 0.33 | -2.69 | 22 | 5.237 | 1.63E-07 | 4.67E-03 |
| cg19123408 | chr16:7104931 | island | NEURL4 | exon+utr5;nc_exon | -0.19 | -4.69 | 22 | -5.173 | 2.30E-07 | 6.08E-03 |
| cg00865429 | chr5:9657286 |  | CLNK | intron | -0.17 | -0.39 | 22 | -5.037 | 4.73E-07 | 1.16E-02 |
| cg06521647 | chr16:68646709 |  | LOC114673004 | intergenic | 0.31 | -5.42 | 22 | 5.02 | 5.18E-07 | 1.19E-02 |
| cg11967527 | chr17:59348130 |  | SCEL | intergenic | -0.75 | 0.45 | 22 | -5.007 | 5.53E-07 | 1.19E-02 |
| cg25212847 | chr11:79884892 |  | OTOGL | cds | -0.33 | 1.45 | 22 | -4.896 | 9.76E-07 | 1.96E-02 |
| cg24913168 | chr17:22950358 |  | SERP2 | exon+utr3 | -0.25 | 4.26 | 22 | -4.88 | 1.06E-06 | 1.96E-02 |
| cg14287486 | chr14:126985171 |  | GLB1L2;LOC114672451 | cds;nc_exon | 0.53 | -4.76 | 22 | 4.876 | 1.08E-06 | 1.96E-02 |
| cg23258615 | chr9:57965324 |  | ANXA11 | intron+utr5 | -0.67 | -2.76 | 22 | -4.846 | 1.26E-06 | 2.10E-02 |
| cg24329497 | chr1:178340010 |  | PIK3R3 | intron | -0.22 | 1.07 | 22 | -4.842 | 1.29E-06 | 2.10E-02 |
| cg15122985 | chr10:99285758 |  | TCEA2 | intron+utr5 | -0.13 | 0.57 | 22 | -4.814 | 1.48E-06 | 2.31E-02 |
| cg23899409 | chr19:54148763 |  | LILRA4 | intergenic | 0.44 | -4.42 | 22 | 4.764 | 1.90E-06 | 2.84E-02 |
| cg08226752 | chr9:118036118 | island | EMX2 | intergenic | -0.22 | -2.53 | 22 | -4.73 | 2.25E-06 | 3.21E-02 |
| cg14477320 | chr19:10310014 | shore | KRI1 | intron | -0.29 | 3.57 | 22 | -4.715 | 2.42E-06 | 3.30E-02 |
| cg11791144 | chr8:32593638 |  | NRG1 | intron | -0.2 | 0.4 | 22 | -4.706 | 2.53E-06 | 3.30E-02 |
| cg09008926 | chr7:34162676 |  | POLR2M | intergenic | -0.49 | 2.51 | 22 | -4.694 | 2.67E-06 | 3.30E-02 |
| cg25867754 | chr2:37453438 |  | EPHB1 | intron | 0.38 | -6.13 | 22 | 4.691 | 2.73E-06 | 3.30E-02 |
| cg16580880 | chr1:205684531 |  | LOC114673978 | intergenic | 0.43 | -0.47 | 22 | 4.686 | 2.78E-06 | 3.30E-02 |
| cg15730274 | chr2:28824160 |  | LOC114676536 | intergenic | -0.45 | 1.61 | 22 | -4.668 | 3.04E-06 | 3.34E-02 |
| cg20154457 | chr11:7410758 | shore | PEX5 | intron | -0.29 | 4.27 | 22 | -4.667 | 3.06E-06 | 3.34E-02 |
| cg03387753 | chr7:17170707 |  | RAD51 | cds | -0.79 | 2.74 | 22 | -4.662 | 3.13E-06 | 3.34E-02 |
| cg20559009 | chr11:124579719 |  | LOC114670987 | intergenic | -0.16 | 0.24 | 22 | -4.651 | 3.30E-06 | 3.34E-02 |
| cg08478189 | chr12:94580677 | shore | KLF7 | exon+utr5;intron+utr5 | -0.27 | -4.9 | 22 | -4.651 | 3.31E-06 | 3.34E-02 |
| cg04145422 | chr15:14987934 |  | MAPKAP1 | intron;intron+utr5 | -0.52 | 2.87 | 22 | -4.644 | 3.42E-06 | 3.35E-02 |
| cg21888978 | chr3:98915266 |  | CRPPA | intergenic | 0.51 | -3.99 | 22 | 4.623 | 3.79E-06 | 3.56E-02 |
| cg21792008 | chr9:129392270 |  | LOC107000309 | intergenic | -0.18 | -0.47 | 22 | -4.615 | 3.93E-06 | 3.56E-02 |
| cg09795075 | chr7:1625165 |  | GABRG3 | intron | 0.39 | -4.44 | 22 | 4.615 | 3.94E-06 | 3.56E-02 |
| cg24501728 | chr1:9363375 |  | CHRM3 | intron+utr5 | -0.79 | 4.85 | 22 | -4.607 | 4.09E-06 | 3.56E-02 |
| cg26631437 | chr8:31135768 |  | TEX15;LOC114680220 | intron;nc_intron | 0.27 | -4.58 | 22 | 4.604 | 4.15E-06 | 3.56E-02 |
| cg18629535 | chr11:112507638 | island | RPH3A | intron+utr5;exon+utr5 | -0.22 | -2.38 | 22 | -4.586 | 4.52E-06 | 3.78E-02 |
| cg17194154 | chr17:76492727 | shore | SOX21 | intron+utr3 | 0.27 | -5.7 | 22 | 4.564 | 5.03E-06 | 4.08E-02 |
| cg02660117 | chr1:130304100 | shore | MIR760 | intergenic | -0.37 | 0.14 | 22 | -4.56 | 5.11E-06 | 4.08E-02 |
| cg00263589 | chr7:86170675 |  | NRL | intron+utr5 | -0.17 | 0.92 | 22 | -4.552 | 5.33E-06 | 4.16E-02 |
| cg02177621 | chr7:114829428 | shore | STYX | intergenic | 0.23 | -5.02 | 22 | 4.519 | 6.21E-06 | 4.66E-02 |
| cg21542081 | chr7:10312568 |  | AVEN | intron | -0.73 | 1.89 | 22 | -4.518 | 6.24E-06 | 4.66E-02 |
| cg00740870 | chr7:129680496 |  | PLEKHH1 | nc_exon;cds | 0.3 | 0.28 | 22 | 4.508 | 6.53E-06 | 4.77E-02 |
| cg05900177 | chr5:73820341 |  | LOC114678529 | intergenic | 0.37 | -3.41 | 22 | 4.501 | 6.76E-06 | 4.84E-02 |
| cg03412153 | chr9:14251637 | shore | OPTN | intergenic | 0.4 | -6.08 | 22 | 4.496 | 6.91E-06 | 4.84E-02 |

